# A rat model of cirrhosis with well-differentiated hepatocellular carcinoma induced by thioacetamide

**DOI:** 10.1101/2024.04.18.590120

**Authors:** Zhiping Hu, Takeshi Kurihara, Yiyue Sun, Zeliha Cetin, Rodrigo M. Florentino, Lanuza A. P. Faccioli, Zhenghao Liu, Bo Yang, Alina Ostrowska, Alejandro Soto-Gutierrez, Evan R. Delgado

## Abstract

Hepatocellular carcinoma (HCC) is a leading cause of cancer-related deaths, and commonly associated with hepatic fibrosis or cirrhosis. This study aims to establish a rat model mimicking the progression from liver fibrosis to cirrhosis and subsequently to HCC using thioacetamide (TAA). We utilized male Lewis rats, treating them with intra-peritoneal injections of TAA. These rats received bi-weekly injections of either 200 mg/kg TAA or saline (as a control) over a period of 34 weeks. The development of cirrhosis and hepatocarcinogenesis was monitored through histopathological examinations, biochemical markers, and immunohistochemical analyses. Our results demonstrated that chronic TAA administration induced cirrhosis and well-differentiated HCC, characterized by increased fibrosis, altered liver architecture, and enhanced hepatocyte proliferation. Biochemical analyses revealed significant alterations in liver function markers, including elevated alpha-fetoprotein (AFP) levels, without affecting kidney function or causing significant weight loss or mortality in rats. This TAA-induced cirrhosis and HCC rat model successfully replicates the clinical progression of human HCC, including liver function impairment and early-stage liver cancer characteristics. It presents a valuable tool for future research on the mechanisms of antitumor drugs in tumor initiation and development.

## Introduction

Liver cancer stands as the second most common cause of death from cancer, with hepatocellular carcinoma (HCC) being the primary type responsible for this high mortality rate^1^. Remarkably, about 90% of HCC cases are closely associated with liver fibrosis or cirrhosis, both of which are outcomes of chronic liver damage. Although various carcinogenic pathways may be involved in these conditions, the transformation of liver tissue into a malignant state is significantly influenced by liver fibrosis or cirrhosis^2^. This transformation process impacts several critical aspects of liver function, including angiogenesis, the composition of the extracellular matrix, and the metabolism of drugs. These changes highlight the critical need for comprehensive animal models that combine features of both cirrhosis and HCC. Such models are essential for evaluating the efficacy of anticancer drugs in preventing or slowing the progression of liver cancer^2,3^.

Current animal models, however, fall short in accurately representing all stages of liver fibrosis, particularly the transition to the cirrhotic stage^4–6^. In addition to not molecularly replicating genetic alterations observed in the clinic, even the widely used diethylnitrosamine-impaired rat (DEN rat) model, though effective in simulating human hepatocarcinogenesis, does not fully replicate the entire progression from liver fibrosis to cirrhosis and eventually to HCC^7–9^. To address these limitations, our study aims to develop a rat model induced by thioacetamide (TAA). This model is designed to closely mimic the progression from liver fibrosis to cirrhosis, ultimately leading to the development of HCC. Utilizing this model, we intend to conduct thorough research and investigations into the effectiveness of antitumor drugs and explore the tumor microenvironment in greater depth.

## Materials and Methods

### Animal models

In this research, male Lewis rats aged 4 weeks and weighing between 100-130 grams were utilized. These rats were sourced from Charles River Laboratories (MA, USA). They were housed in pairs in isolated cages within the Department of Laboratory and Animal Resources at the University of Pittsburgh. The environment for the animals was controlled for temperature and light/dark cycles. The rats had access to a standard diet and water.

The experimental procedure involved administering intra-peritoneal injections of TAA at a dosage of 200 mg/kg, sourced from Sigma Chemical Co. (St. Louis, MO, USA). For the control group, saline was used. These injections were given twice weekly, with the TAA being dissolved in a saline solution at a concentration of 100mg/mL.

Blood samples were collected from both the TAA-treated and saline-treated (control) rats at four distinct time points: prior to the start of treatment (Week 0), and at Weeks 12, 26, and 34. Tissue samples for analysis were collected at Week 34 from the control group, which received saline injections, and at Weeks 26 and 34 from the TAA-treated group.

This study adhered strictly to the Guiding Principles for the Care and Use of Laboratory Animals as established by the University of Pittsburgh. The experimental protocol received approval from the Animal Care Ethics Committee and underwent review by the local ethics committee, ensuring compliance with ethical standards in animal research.

### Chemical parameters tests

0.5mL blood samples for each rat were collected by puncture of the tail vein. Samples were analyzed by Zoetis Abaxis VetScan VS2 (Abaxis, Union City, CA) with Preventive Care Profile Plus Rotor was used to quantitatively measure the following variables: albumin(ALB), alanine aminotransferase(ALT), alkaline phosphatase(ALP), aspartate aminotransferase (AST), blood urea nitrogen(BUN), calcium(CA), chloride(CL-), creatine(CRE), globulin(GLOB), glucose (GLU), potassium (K+), sodium (NA+), total carbon dioxide(tCO2), total bilirubin(TBIL), total protein(TP). INR was analyzed by CoaguChek® XS system (Roche diagnostics). AFP was analyzed by Rat αFP(Alpha-Fetoprotein) ELISA Kit (Elabscience).

### Tissue section analyses

The liver tissues were fixed in 4% paraformaldehyde (PFA) and then paraffin-embedded; four-micrometer sections of tissue were prepared. Hematoxylin-eosin (HE) staining was used for the histopathological examination. Sirius red staining according to the manufacturer’s protocols (Sigma-Aldrich) was used to detect the fibrosis development. To ascertain the HCC development, the immunohistochemical examination of localization of GST-P protein and Glypican 3 protein was performed perceptively in liver sections. To detect vascularization, immunohistochemical examination of localization of CD34 protein was performed in liver sections. To assess liver architecture to show the thickness of hepatocyte plates, the special staining of reticulin fibers (type III collagen) in the space of Disse was performed. To detect proliferating cells, immunohistochemical examination of the Ki67 was performed. For all the immunohistochemical examination, 4% paraformaldehyde-fixed liver sections were applied by using the avidin-biotin complex method. Briefly, after deparaffinization (and target retrieval using Target Retrieval Citrate buffer Solution in microwave in the cases of Glypican 3, CD34, Ki67), the sections were treated sequentially with 3% H2O2, normal goat serum in the cases of Ki67 and GST-P or normal or horse serum in the cases of CD 34 and Glypican 3, primary antibody( ie, mouse anti-rat monoclonal IgG1 Glypican 3(,1:200, room temperature, 1 hour, ab216606/Abcam), rabbit anti-rat monoclonal IgG CD34(1:100, room temperature, 1 hour, ab81289/abcam), mouse anti-rat monoclonal IgG Ki67(1:50, room temperature, 1 hour, 550609/BD ) and rabbit anti-rat GST-P polyclonal antibody (ready to use, room temperature, 1 hour; 311-H/MBL), biotin-labeled goat anti-rabbit IgG or biotin-labeled horse anti-mouse IgG and avidin-biotin-peroxidase complex (VECTASTAIN Elite ABC HRP kit; Vector Laboratories PK6101). Counterstaining by hematoxylin for 30 seconds. Images were captured by using the Olympus IX71 inverted microscope (Olympus, Tokyo, Japan) and collected by cellSens Dimension software. The positive area threshold was quantified using ImageJ software (NIH, Bethesda, MD, USA).

## Statistical Analysis

All the data were tested for normality and the appropriate statistical test was chosen. The comparisons of means were calculated by using ANOVA tests with Tukey HSD correction for multiple means comparisons, and independent T-tests only when two means were compared. The data are presented as mean values _ standard error mean (SEM). The statistical analyses were performed using Prism 10 (GraphPad Software Inc., San Diego, CA, USA).

## Result

### Characterization of Chronic TAA-induced Liver Cirrhosis

To delineate the progression of cirrhosis and hepatocarcinogenesis induced by TAA, a total of 32 Lewis rats were subjected to a regimen of bi-weekly intra-peritoneal injections. The control group received saline injections for 34 weeks(n=16), while the experimental group was administered TAA at a dosage of 200 mg/kg for two distinct durations: 26 weeks(n=8) and 34 weeks(n=8) (Figure 1A). Notably, TAA treatment inhibited body weight gain in the rats (Figure 1B) and induced progressive liver damage, culminating in the development of cirrhotic nodules in 100% of the TAA-treated animals after 26 weeks of injections (Figures 1C and 1D). To assess the extent of fibrosis, Sirius red staining was employed. This histological analysis revealed a significant increase in fibrosis in the TAA-treated rats at both the 26-week (p = 0.0009) and 34-week (p < 0.0001) time points, compared to the control group (ANOVA, p < 0.0001) (Figure 1E). These results demonstrate the efficacy of TAA in inducing liver cirrhosis in a controlled experimental setting, providing a valuable model for studying the pathophysiological mechanisms underlying liver disease progression.

**Figure 1:**
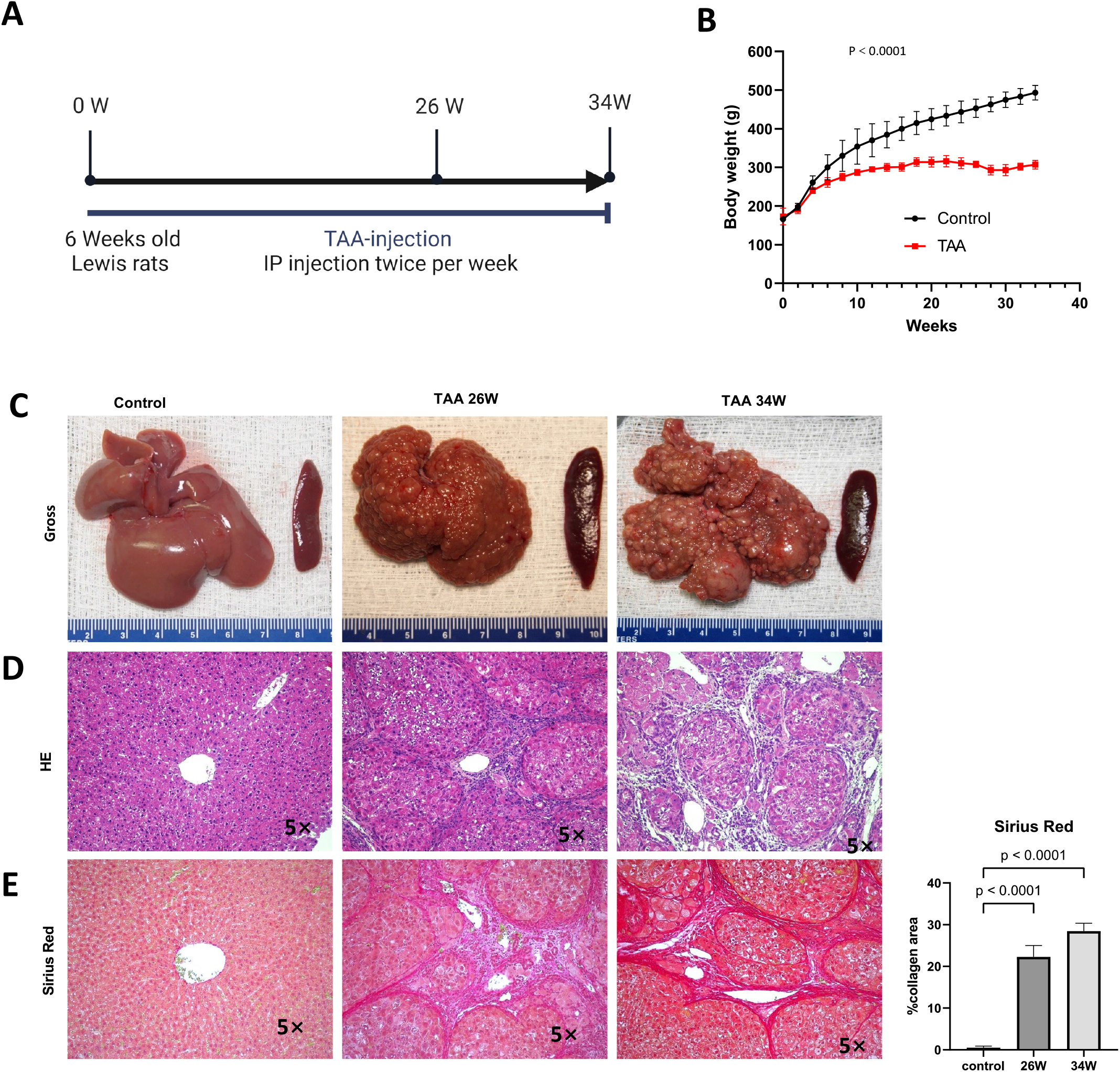
Chronic TAA-Induced Cirrhosis. **A:** Timeline of the protocol for the TAA-induced cirrhotic rat model of HCC. Six-week-old Lewis rats received bi-weekly injections (200 mg/kg). **B:** Graphical representation of rat body weight change in both TAA injection group and saline injection group during treatment. The x-axis represents the treatment duration rather than rat age. Statistics evaluation was performed via two-way ANOVA. p<0.0001, n=15 (Control group), n=15 (TAA group) **C:** Photographic documentation of rat liver morphology and histological staining images (H&E, Sirius Red) at 26, 34 weeks after TAA treatment. Saline injection group used as control. Histological sections were obtained from rat tissue using paraffin embedding techniques. Magnification 5x. Graph on right representing Sirius staining quantification in C by identifying Sirius red-stained area per high power field (HPF). Statistic analysis was performed by two-way ANOVA following Šídák’s multiple comparisons test between control group and two time points in TAA group. Sirius red stained area in two TAA group is significantly higher than control group (Both p<0.0001, n=5).

### Phenotypic Characterization of Chronic TAA-Induced Hepatocarcinogenesis HCC

To investigate the hepatocarcinogenic effects of chronic TAA administration, we focused on cellular differentiation and proliferation. The presence of GST-P positive lesions, indicative of preneoplastic and neoplastic changes, was significantly evident following 26 weeks of TAA treatment (p < 0.0001), with a notable expansion observed at 34 weeks (p < 0.0001) (Figure 2A). Hematoxylin and eosin (HE) staining of GST-P positive lesions in the liver tissues at 26 weeks post-TAA treatment revealed characteristics of cirrhotic structures. By 34 weeks, the lesions exhibited features typical of well-differentiated HCC, including enriched hepatocyte cytoplasm, increased nucleoplasm ratio, and expanded hepatocyte plates (Figure 2B). Further histological analysis involved reticulin and CD34 staining to assess liver architecture and vascular organization. In control rats, the reticulin network was well-preserved. In contrast, TAA-treated rats at 26 weeks displayed a normal reticulin pattern with thick expression along the liver sinusoids and hepatic trabeculae, involving fewer than three cell layers. However, by 34 weeks of TAA treatment, a marked disappearance and collapse of the reticulin network were observed (Figure 2C). Additionally, CD34 staining at 34 weeks post-TAA treatment showed diffused positivity compared to controls and the 26-week TAA group (Figure 2D). These findings align with established criteria for diagnosing well-differentiated HCC, where the absence or reduction of reticulin stain and abnormal reticulin patterns with widened trabeculae are considered reliable indicators (Hong, Patonay, & Finley, 2011). Moreover, Glypican-3 staining, which is associated with advanced HCC stages (Takahashi et al., 2021), was negative in the lesions at 34 weeks of TAA treatment (n=6) (Figure 2E), suggesting a well-differentiated state of HCC. The proliferative capacity of hepatocytes was assessed using Ki67 staining, which revealed a significant increase in Ki67 positive cells at 34 weeks of TAA treatment compared to both the control group and the 26-week TAA group (p < 0.0001) (Figure 3F). These results collectively indicate that chronic TAA administration leads to the development of well-differentiated HCC, characterized by specific histopathological changes and increased hepatocyte proliferation.

**Figure 2:**
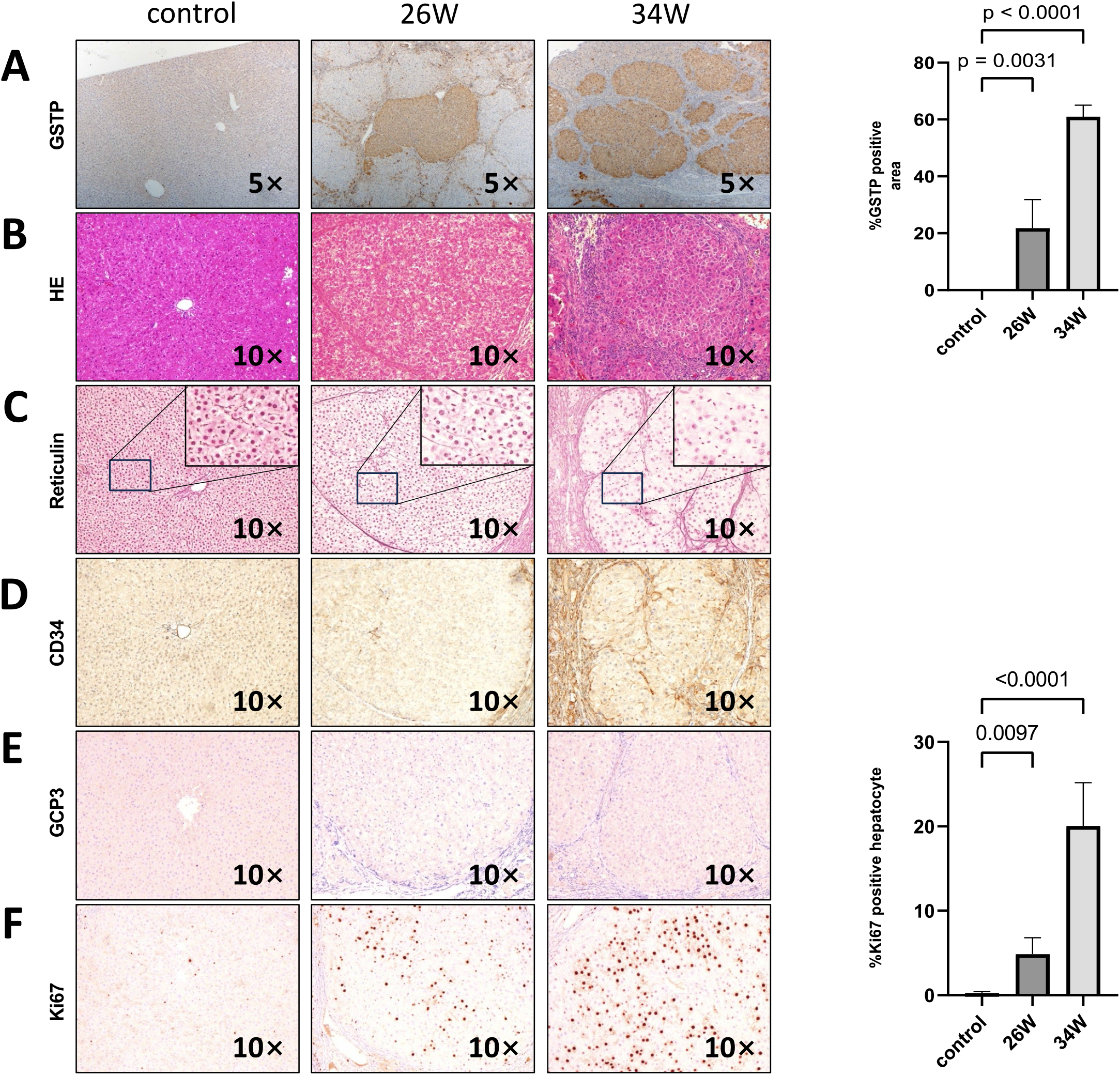
Chronic TAA-Induced Hepatocarcinogenesis. Photographic documentation of rat liver histological staining images and immunohistochemistry (IHC) staining images at 26 and 34 weeks after TAA treatment. Saline injection group used as control. Rat liver sections were obtained using paraffin embedding techniques. **(A)** GST-P IHC staining, with GST-P positive quantification of GST-P positive nodule surface area per HPF on the right side. Statistics analysis was performed via two-way ANOVA following Šídák’s multiple comparisons test between control group and two time points in TAA group. GST-P positive rate in two TAA group is significantly higher compared to control group(26W: p=0.0031, n=5, 34W: p<0.0001, n=5). Magnification 5x. **(B)** H&E staining, magnification 10x. **(C)** Reticulin staining, magnification 40x. **(D)** CD34 IHC staining, magnification 10x. **(E)** GCP3 IHC staining, magnification 10x. **(F)** Ki67 IHC staining, with quantification of Ki67 positive nodule numbers and surface area per HPF on the right side. Statistic analysis method same with A, Ki67 positive rate in two TAA group is significantly higher compared to control group(26W: p=0.0097, n=5, 34W: p<0.0001, n=5)

**Figure 3:**
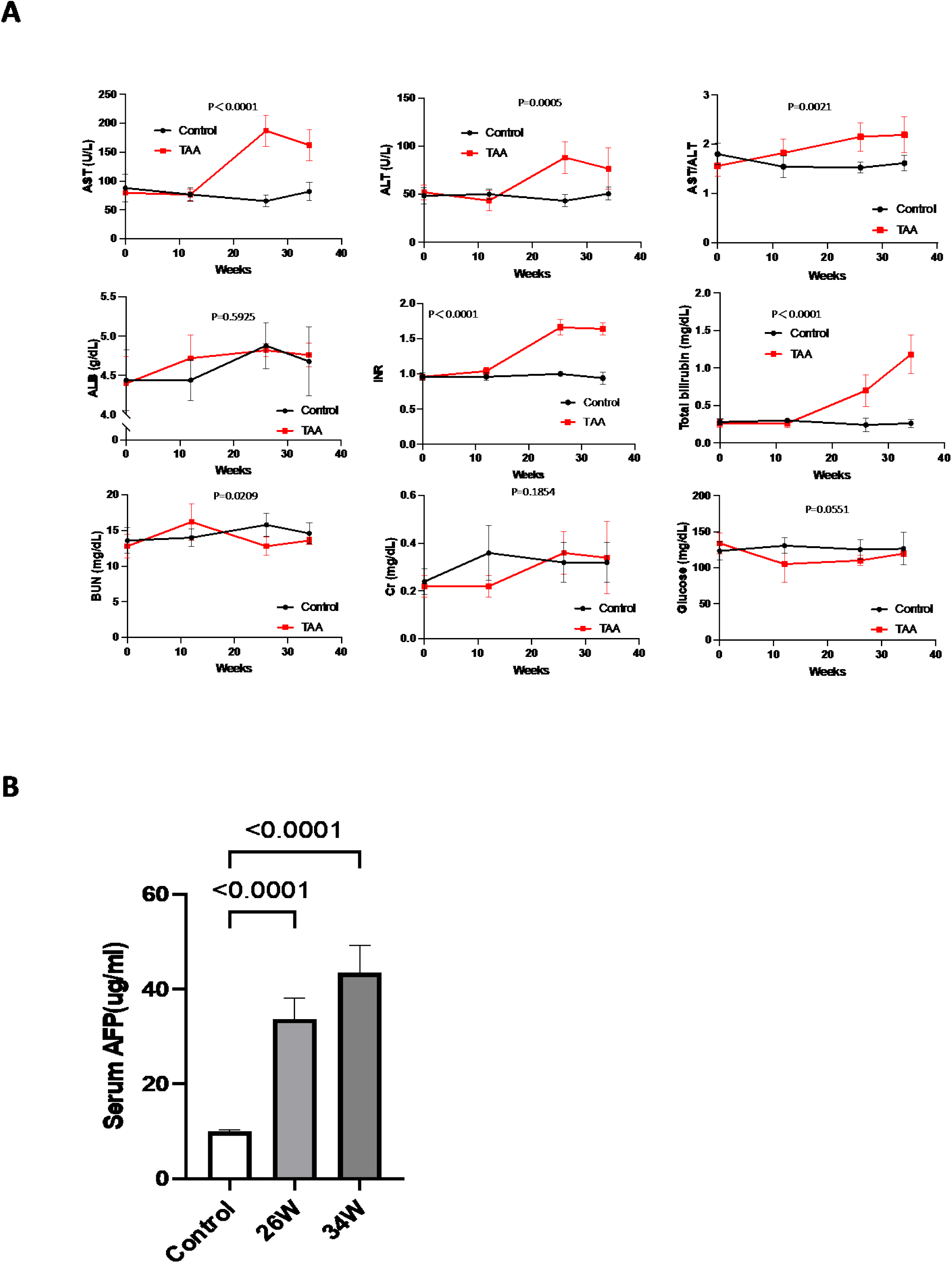
Biochemical Markers, Body Weight Changes, and Serum AFP Levels During Chronic TAA-Induced Hepatocarcinogenesis. A: Graphic representation of variations in clinically-relevant serum parameters for rats in both the TAA injection group and the saline injection control group during treatment, p values showed in graph. Statistics analyses were performed using two-way ANOVA. Post-TAA treatment, a significant increase in AST, ALT, AST/ALT ratio, PT/INR and total bilirubin level was observed while other parameters tested remains no significant change. B: Graphic representation of serum AFP levels in control, 26-week, and 34-week TAA-treated groups by ELISA. Rat serum was isolated from fresh blood samples by centrifugation. Statistics analysis was performed using two-way ANOVA following Šídák’s multiple comparisons test between control group and two time points in TAA group. Post-TAA treatment, a significant increase in serum AFP level was observed (both p<0.0001, n=8)

### Biochemical Profile of Animals Under Chronic TAA-Induced Hepatocarcinogenesis in cirrhotic liver

Biochemical analysis revealed notable changes in the TAA-induced hepatocarcinogenesis model. Specifically, levels of ALT, AST, the ALT/AST ratio, TBIL, and INR were significantly elevated in the TAA group compared to the control group at both week 26 and week 34 (Figure 3A). In contrast, levels of ALB, BUN, Cr, and glucose did not show any significant difference between the TAA-treated group and the control group (Figure 3A). Additionally, the level of serum AFP, a marker often associated with liver cancer, was found to be higher in the TAA group at both week 26 and week 34 compared to the control group (Figure 3B). This increase in AFP levels further corroborates the development of hepatocarcinogenesis in the TAA-treated rats.

Based on these results, a thioacetamide-induced cirrhosis combined with HCC rat model was established, exhibiting HCC pathological characteristics such as hepatocyte proliferation, liver fibrosis/cirrhosis, widened hepatocyte plates, and disorganized vasculature. Additionally, it displayed parameters of chronic liver function impairment, thereby replicating the typical processes observed in the progression of human HCC.

## Discussion

The significant association of HCC with chronic liver damage, leading to fibrosis or cirrhosis, is underscored by the fact that approximately 90% of HCC cases arise from these conditions. Liver fibrosis or cirrhosis plays a crucial role in the malignant transformation of liver tissue, as highlighted in various studies^10^. The changes induced by these conditions, such as altered liver vascular formation, modifications in the extracellular matrix composition, and impacts on drug metabolism, emphasize the necessity of employing animal models that effectively combine features of both cirrhosis and liver cancer. These models are instrumental in investigating the mechanisms through which anti-tumor drugs influence tumor initiation and development^4,11,12^.

Current rodent models, particularly mouse models, have limitations in fully replicating all stages of liver fibrosis, especially the transition to cirrhosis^4–6^. The diethylnitrosamine (DEN) induced rat model, commonly used to simulate human HCC in a relatively short time injection, does not completely mimic the progression from liver fibrosis to cirrhosis and subsequent HCC development^6–8^. In response to this gap, our study developed a TAA-induced rat model that effectively combines cirrhosis with HCC, closely mirroring the progression observed in human HCC. This model, through specific staining and histological examination, predominantly exhibits well-differentiated HCC, aligning with the increasing prevalence of early-stage liver cancers in clinical settings.

Biochemically, the TAA-induced cirrhosis and HCC model demonstrated significant elevations in AFP levels, a key tumor marker for HCC, suggesting the onset of liver carcinogenesis^1,13,14^. Other indicators, such as increased INR, elevated TBIL levels, high AST and relatively lower ALT levels, pointed towards abnormal liver function. Notably, the AST/ALT ratio greater than 2, typically seen in alcoholic liver cirrhosis, was also observed in our model, suggesting potential mitochondrial damage caused by TAA metabolites^15^. The elevated INR is indicative of impaired liver synthetic function, the increased TBIL could be attributed to bilirubin metabolism dysfunction. However, the maintenance of normal albumin and blood sugar levels, along with the absence of ascites and hepatic encephalopathy, indicated that liver impairment was not end-stage. Additionally, normal kidney function suggested that TAA did not induce renal damage. Compared to other rodent models of HCC^2,5,7,8^, our model demonstrated liver function impairment without affecting other organs, thereby more closely resembling the clinical characteristics of HCC.

Nevertheless, this model possesses certain limitations. To achieve a 100% incidence rate of HCC, a prolonged period exceeding 34 weeks was necessary. Furthermore, additional research is required to elucidate the molecular mechanisms underpinning hepatocarcinogenesis associated with TAA-induced cirrhosis.

In conclusion, the establishment of a rat model with TAA-induced cirrhosis and HCC in this study offers a comprehensive understanding of the pathological characteristics and progression of early-stage liver cancer with concurrent liver function impairment. This model provides a more accurate reflection of clinical features than other animal models and is crucial for future research into the mechanisms of anti-tumor drugs in tumor initiation and progression.

## Authors’ Contributions

Conceived and designed the study: H.Z., T.K., R.F., A.S-G.

Performed data acquisition: H.Z., T.K., Y.S., Z.C., B.Y., A.S-G.

Analyzed and interpreted data: H.Z., T.K., Y.S., B.Y., R.F., L.F., A.O., A.S-G.

Writing, review, and/or revision of the manuscript: H.Z., T.K., A.S-G., E.R.D.

## Financial support

E.R.D.: K22 CA258677

A.S-G.: R01 DK099257, UH3 TR003289, P01 DK096990, R01 DK117881, UH3 DK119973, U01 TR002383

Human Synthetic Liver Biology Core and the Pittsburgh Liver Research Center: 1P30DK120531-01

Pittsburgh Liver Research Center: NIH/NIDDK P30DK120531

Center for Transcriptional Medicine at the University of Pittsburgh

